# TWISTED DWARF1 regulates *Arabidopsis* stamen development by differential activation of ABCB-mediated auxin transport

**DOI:** 10.1101/2021.11.30.470611

**Authors:** Jie Liu, Roberta Ghelli, Maura Cardarelli, Markus Geisler

## Abstract

Despite clear evidence that a local accumulation of auxin is likewise critical for male fertility, much less is known about the components that regulate auxin-controlled stamen development.

In this study, we analyzed physiological and morphological parameters in mutants of key players of ABCB-mediated auxin transport and spatially and temporally dissected their expression on the protein level as well as auxin fluxes in the Arabidopsis stamens. Our analyses revealed that the FKBP42, TWISTED DWARF1 (TWD1), promotes stamen elongation and, to a lesser extent, anther dehiscence, as well as pollen maturation and thus is required for seed development. Most of the described developmental defects in *twd1* are shared with the *abcb1 abcb19* mutant, which can be attributed to the fact that TWD1 - as a described ABCB chaperon - is a positive regulator of ABCB1 and ABCB19-mediated auxin transport. However, reduced stamen number was dependent on TWD1 but not on investigated ABCBs, suggesting additional actors down-stream of TWD1. We predict an overall housekeeping function for ABCB1 during earlier stages, while ABCB19 seems to be responsible for the key event of rapid elongation at later stages of stamen development. Our data indicate that TWD1 controls stamen development by differential activation of ABCB-mediated auxin transport in the stamen.

**Highlight:** By using a mix of phenotypical and imaging analyses, we here identify and functionally characterize a new master regulator of flower development.

## Introduction

Auxin is an essential hormone that is implicated in virtually all aspects of plant development, including plant organogenesis and pattern formation (Woodward and Bartel, 2005). The finding that plants cultured in the presence of auxin transport inhibitors, such as 1-*N*-naphtylphtalamic acid (NPA) (Okada et al., 1991; Reinhardt et al., 2000) or 2,3,5-triiodobenzoic acid (TIBA) (Thomson et al., 1973), reveal a reduced number of stamens, as well as no floral buds or flowers with no stamens, connected flower development directly to auxin transport. Since the characterization of the auxin transport mutant *pin-formed1* (*pin1*) (Gälweiler et al., 1998; Okada et al., 1991), auxin is considered a key regulator of floral development. On the other hand, the flower is the most complex structure in plants, making it an excellent but also challenging model for elucidating the molecular mechanisms by which auxin regulates plant growth and development.

The *Arabidopsis thaliana* flower consists of four concentric basic structures: sepals, petals, stamens and carpels. It contains four long stamens and two short stamens, which each consist of two parts, the filament and the anther. The filament has a single vascular bundle that serves as a conduit for water and nutrients, while the anther locules contain the microsporangium that forms the pollen (Sanders et al., 1999). The transverse section of a mature anther has four distinct structures surrounding the anther locule from the inner to the outer: the tapetum, the middle layer, the endothecium and the epidermis (see Supporting Information Fig. S5; (Goldberg et al., 1993; Scott et al., 2004). The entire process of stamen developmental can be divided into an early and a late phase. Stamen morphogenesis occurs at an early stage (stages 5-9 of flower development), followed by the late stages (stages 10-13), consisting of pollen maturation, anther dehiscence and stamen filament elongation (Cecchetti et al., 2008; Feng and Dickinson, 2010; Goldberg *et al*., 1993). During late stamen development, filament elongation is mainly due to cell expansion and is particularly rapid from stages 12 to 13 (Bowman,1994). Very recently, by means of live imaging an atlas of cellular growth dynamics of the stamen was provided (Silveira et al., 2021). It revealed an early specification of the filament and the anther. Formation of the anther lobes is associated with a temporal increase of growth at the lobe surface that correlates with intensive growth of the developing locule (Silveira *et al*., 2021).

Previous studies have provided increasing evidence that an auxin maximum provided by polar auxin transport (PAT) is necessary for both early and late stages of stamen development (Cecchetti et al., 2015). Imaging of the synthetic auxin response reporters, DR5::GFP and DII::Venus, at consecutive time points during flower development revealed that auxin response maxima in the floral meristem colocalize with the positions of organ initiation (Goldental-Cohen et al., 2017; van Mourik et al., 2012). Precise spatiotemporal local auxin biosynthesis and its polar transport are essential for floral meristem initiation and subsequent development (Alvarez-Buylla et al., 2010; Yadav et al., 2020).

The *pin1* mutant fails widely to form lateral flowers (Gälweiler *et al*., 1998; Okada *et al*., 1991). The occasional *pin1* flowers lack sepals and stamens and have wide petals, pistil-like structure with no ovules in the ovary, while exogenous auxin application can restore flower formation on *pin1* inflorescences (Gälweiler et al., 1998; Okada et al., 1991; Reinhardt et al., 2000). In analogy, a defective AGC protein kinase, PINOID, which is thought to regulate both PIN1 polarity and activity by PIN1 phosphorylation (Hammes et al., 2021), results likewise in pin-shaped inflorescence with excessive number of petals, few sepals and stamens (Bennett et al., 1995). Interestingly, genetic *PIN1* or *PINOID* loss-of function can be also phenocopied pharmacologically by treatment with the non-competitive auxin export inhibitor, NPA, suggesting that NPA either inhibits PIN1 activity (Abas et al., 2021) or PIN1 polarity (Friml et al., 2004). Flowers that rarely form stamens or flowers with a reduced number of stamens have also been observed in weaker *PIN1* alleles (e.g. *pin1-3, pin1-4*, and *pin1-5*) (Bennett *et al*., 1995), while *pin3 pin7* double mutant flowers form no stamens at all (Benkova et al., 2003).

Loss-of-function mutation of members of the ATP-binding cassette subfamily B (ABCB) that act also as auxin transporters affects stamen elongation but does not alter the number of stamens (Cecchetti *et al*., 2008; Cecchetti *et al*., 2015; Noh et al., 2001). Cecchetti et al. (2008) showed that *abcb1 abcb19* (formerly referred to as *pgp1 pgp19*) flowers, in addition to short stamen filaments, also exhibit a small percentage of early dehiscent anthers. On the other hand, treatment of flower buds at floral stage 11 with NPA resulted in 15–20% of indehiscent anthers at stage 13, suggesting that auxin transport through ABCB1 and ABCB19 regulates the timing of anther dehiscence (Cecchetti et al., 2008).

Interestingly, while the *pin1 abcb19* double mutant shows a single pin-formed inflorescence, the triple *pin1abcb1abcb19* mutant revealed a partial rescue of the *pin1* mutant phenotype and formed a few flowers, suggesting that the loss of *ABCB1* is epistatic to *PIN1* in the floral meristematic tissue. In summary, this indicates that PINs and ABCBs function in the Arabidopsis flower interactively in a tissue-specific manner (Blakeslee et al., 2007). In summary it appears that different PINs are involved in the early phase of flower development, whereas members of both PIN and ABCBs are required during the late developmental phase (Cardarelli and Cecchetti, 2014).

The FKBP42, TWISTED DWARF1 (TWD1)/ULTRACURVATA2 (UCU2), acts as a chaperone during early ABCB biogenesis (Geisler and Hegedus, 2020). As a result, ABCB1,4,19 are widely retained on the endoplasmic reticulum (ER) in the *twd1* mutant, get degraded and thus do rarely reach the plasma membrane (Wang et al., 2013; Wu et al., 2010). As a consequence, *TWD1/UCU2* loss-of-function results in a strong pleiotropic, development syndrome, including dwarfism, a delayed life cycle and a helical, non-handed disorientation of most plant organs and epidermal layers Their flowers are small, and their pedicels, stamens, pistils and siliques are twisted, and fertility is reduced in these mutants (Bailly *et al*., 2006; Geisler et al., 2003; Kamphausen et al., 2002; Perez-Perez et al., 2004). This phenotype is very similar (but not identical) to the *abcb1abcb19* mutant (Bailly *et al*., 2006; Geisler and Bailly, 2007; Geisler *et al*., 2003; Wang *et al*., 2013; Wu *et al*., 2010; Zhu et al., 2016). However, the impact of TWD1 on individual ABCB proteins in regulating the timing of anther dehiscence in these processes is still elusive.

Here, by analyzing physiological and morphological parameters and auxin fluxes in the *twd1* flower, we demonstrate an essential role for TWD1 in the regulation of male gametophyte development. Our data indicate that TWD1 controls stamen development by differential activation of ABCB-mediated auxin transport.

## Materials and Methods

### Accession numbers

Sequence data from this article can be found in the Arabidopsis Genome Initiative or GenBank/EMBL databases under the following accession numbers: *TWD1*, At3g21640; *ABCB1* (formerly *PGP1*), At2g36910; *ABCB19* (formerly *MDR1* or *PGP19*), At3g28860; *ABCB4* (formerly *MDR4* or PGP4), At2g47000; *PIN2*, At5g57090 and *PIN8*, At5g15100.

### Growth conditions and sample collection

All Arabidopsis plants were grown for 15 days at 22°C under an 8h light/16h dark cycle and then transferred to a 16h light/8h dark cycle until flowering; light intensities were 100 micro mol m^-2^ s^-1^. Anthers or stamens were collected for measurement and classified into four categories according to their stage, as described above. Throughout this study the ecotype Columbia (Col-0) as used and mutant lines are as follow: *twd1* (*twd1-3*, (Geisler *et al*., 2003)), HA-TWD1 (*35S:HA-TWD1* in *twd1-3;* (Geisler et al., 2003)), *abcb1 (abcb1-100), abcb19 (abcb19-101), abcb1 abcb19 (abcb1-100 abcb19-101)*, (Jenness et al., 2020)), *pin2* (*eir1-4*, (Luschnig et al., 1998) and *pin8* (*pin8-1;* (Ding et al., 2012).

### Pollen microscopy

Pollen viability was estimated from flower buds harvested just before anthesis. For each plant, anthers were dissected and observed under a light microscope after Alexander staining (Alexander, 1969). The cytoplasm of viable pollen grains was colored in red, and the pollen wall was colored in green, thus dead pollen grains appeared greenish. Nuclei in microspores were stained with the dye DAPI according to a published protocol (Ross et al., 1996). Developing pollen grains from wild-type and mutant flowers were stained for callose with aniline blue solution (0.05% [w/v] in 100 mM potassium phosphate buffer, pH 8.5) for 5 min (Park and Twell, 2001). Stained pollen grains were observed using a microscope under UV light (DM4000B, Leica).

For confocal laser scanning microscopy work, the Leica TCS-SP5 was used. Various confocal settings were set to record the emission of GFP (excitation 488 nm; emission 505 to 530 nm), CFP (excitation 458 nm; emission 465 to 500 nm), chloroplast autofluorescence (excitation 488 nm; emission 650 to 710 nm). Images were acquired with the ImageJ software (http://imagej.nih.gov/ij/) using identical settings for all samples.

### Morphological, histological and cytological analyses

Starch staining was performed as described by (Ruhlmann et al., 2010). Freshly collected flowers were dipped into Lugol’s iodine solution for 5 min followed by a 1-min vacuum infiltration. The flowers were then rinsed twice in deionized water and then imaged under a dissecting microscope.

### *In Situ* hybridization

A partial TWD1 fragment of 150 bp covering the TPR domain and a part of the calmodulin binding domains (aa 264-314) was PCR amplified and cloned into pGEM-T-Easy Vector (Promega Corp.) and verified by sequencing. Digoxigenin-labeled RNA probes were synthesized by *in-vitro* transcription using the DIG RNA labeling kit (Roche). *In situ* analysis was performed as described previously (Cecchetti et al., 2007; Lopez-Dee et al., 1999). Hybridizations with a TWD1 sense probe is shown in Supplemental Figure 2G.

### Statistical analysis

All statistical analyses were performed using Prism 9.2 (GraphPad Software). Confocal images were processed in Adobe Photoshop 2020 (Adobe Systems) and quantified using ImageJ software (http://imagej.nih.gov/ij/).

## Results

### TWD1 plays a key role in stamen development

TWD1 functions primarily as an ABCB chaperone during endoplasmic reticulum to plasma membrane trafficking, which is required for auxin-mediated cell elongation. Both *twd1-3* and *abcb1abcb19* reveal similar phenotypes in respect to their rosette leaves, roots, and hypocotyls, however, the *twd1* flower morphology has not yet been analyzed in detail (Bailly *et al*., 2006; Geisler and Bailly, 2007; Geisler *et al*., 2003; Wang *et al*., 2013; Wu *et al*., 2010; Zhu *et al*., 2016). In contrast, previous studies have shown that the *abcb1 abcb19* double mutant has poor flower fertility because the stamen filaments were 20% shorter than those of the wild type and thus were not sufficiently elongated to position the anthers above the stigma during anthesis (Cecchetti et al., 2008; Noh et al., 2001).

The flower morphology of *twd1-3* is characterized by fewer and smaller flowers compared with the wild type (Figure 1A and Figure 2A). At flowering stage 13, 55% of *twd1-3* flowers were morphologically similar to wild-type flowers, only smaller in size. However, 35% of *twd1-3* flowers showed a curly phenotype (Supporting Information Fig. S1E). Moreover, the petals of *twd1-3* flowers failed to fully expand, and 10% of flowers had only three petals (Figure 1A; Supporting Information Fig. S1E). After careful removal of the petals and sepals, we found that the *twd1-3* mutation affects also the physical position of the anthers. The ratio between stamen and pistil lengths defines the relative position between the anther and the stigma. We discovered that the stamen/pistil ratio is significantly lower at anthesis in *twd1-3* (0.822 ± 0.007) compared to wild-type flowers (1.051 ± 0.005) and comparative to *abcb1abcb19 (0.778* ± 0.005), suggesting that *twd1* stamen filaments are not sufficiently elongated during the pollination phase to position the anthers above the stigma (Figure 1B; Supporting Information Fig. S1B). Microscopic analysis showed that the length of epidermal cells in the middle of *twd* (138.3 ± 1.7 μm) and *abcb1abcb19* stamen (142.0 ± 2.2 μm) was significantly reduced compared to the wild type (194.8 ± 2.9 μm), while the cell width was significantly increased (wild type, 11.8 ± 0.13 μm; *twd1*, 14.1 ± 0.1 μm; *abcb1abcb19*, 13.9 ± 0.1 μm) (Figure 1D-F; Supporting Information Fig. S1C-E). The above described phenotypes of the *twd1* mutant showed very high similarity to those of the *abcb1abcb19* double mutant and can be rescued by expressing *35S:HA-TWD1* in the *twd1-3* background (Geisler *et al*., 2003). This suggests that TWD1 is involved in regulating the rapid elongation of long stamen filaments at stage 12 by a mechanism that may be similar to that in the root and the hypocotyl, where TWD1 is involved in auxin-mediated cell elongation provided by ABCBs (Bailly et al., 2014; Wang *et al*., 2013).

**Fig. 1:**
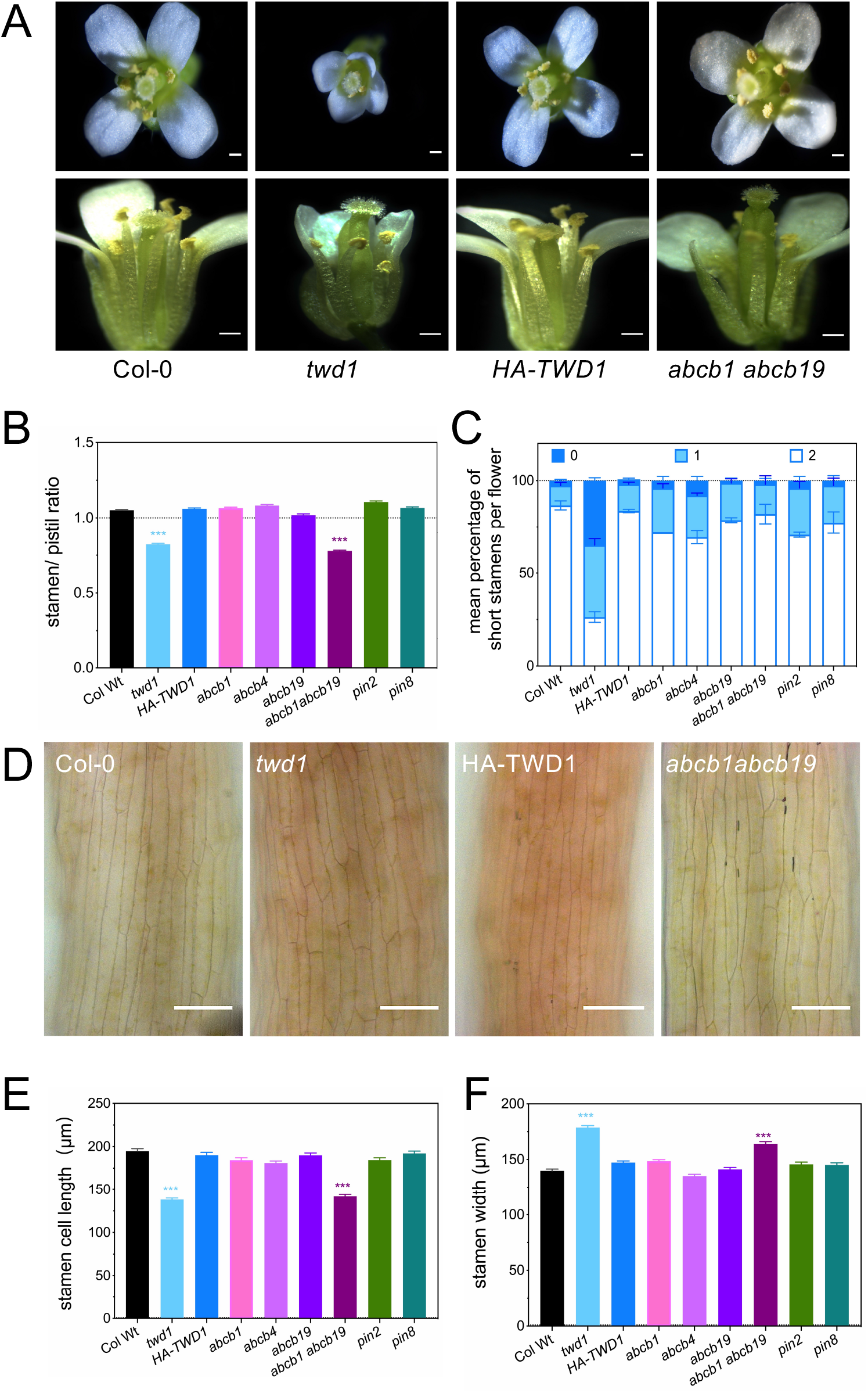
*TWD1* loss-of-function flowers have a defect in stamen development. **A-C:** Flowers of wild-type (Col-0), *twd1 (twd1-3)*, HA-TWD1 (*35S:HA-TWD1* in *twd1-3)* and *abcb1 abcb19 (abcb1-100 abcb19-101)* from flowers at floral stage 13 were imaged (**A**) and stamen/pistil ratios of long stamen (**B**) and the percentage of short stamen (**C**) were quantified. Note that *twd1* and *abcb1 abcb19* show shorter stamen, while *twd1* but not *abcb1 abcb19* shows defects in short stamen development. Bars, 500 μm. Significant differences of mean values ± SEM (ordinary one-way ANOVA, Kruskal-Wallis test) to wild-type are indicated by asterisks (*, p<0.05; **, p<0.01; *** p<0.001). One hundred flowers from four individual plants of each genotype were analyzed. **D-F:** The epidermis in the middle region of wild-type (Col-0), *twd1 (twd1-3), HA-TWD1 (35S:HA-TWD1* in *twd1-3) abcb1 abcb19 (abcb1-100 abcb19-101), pin2 (eir1-4)* and *pin8 (pin8-1)* stamen filaments from flowers at stage 13 was imaged (**D**) and cell length (**E**) and width (**F**) was quantified. Significant differences of mean values ± SEM (ordinary one-way ANOVA, Kruskal-Wallis test) to wild-type are indicated by asterisks (*, p<0.05; **, p<0.01; *** p<0.001). One hundred flowers from four individual plants of each genotype were randomly selected and measured. Bars, 50 μm.

**Fig. 2:**
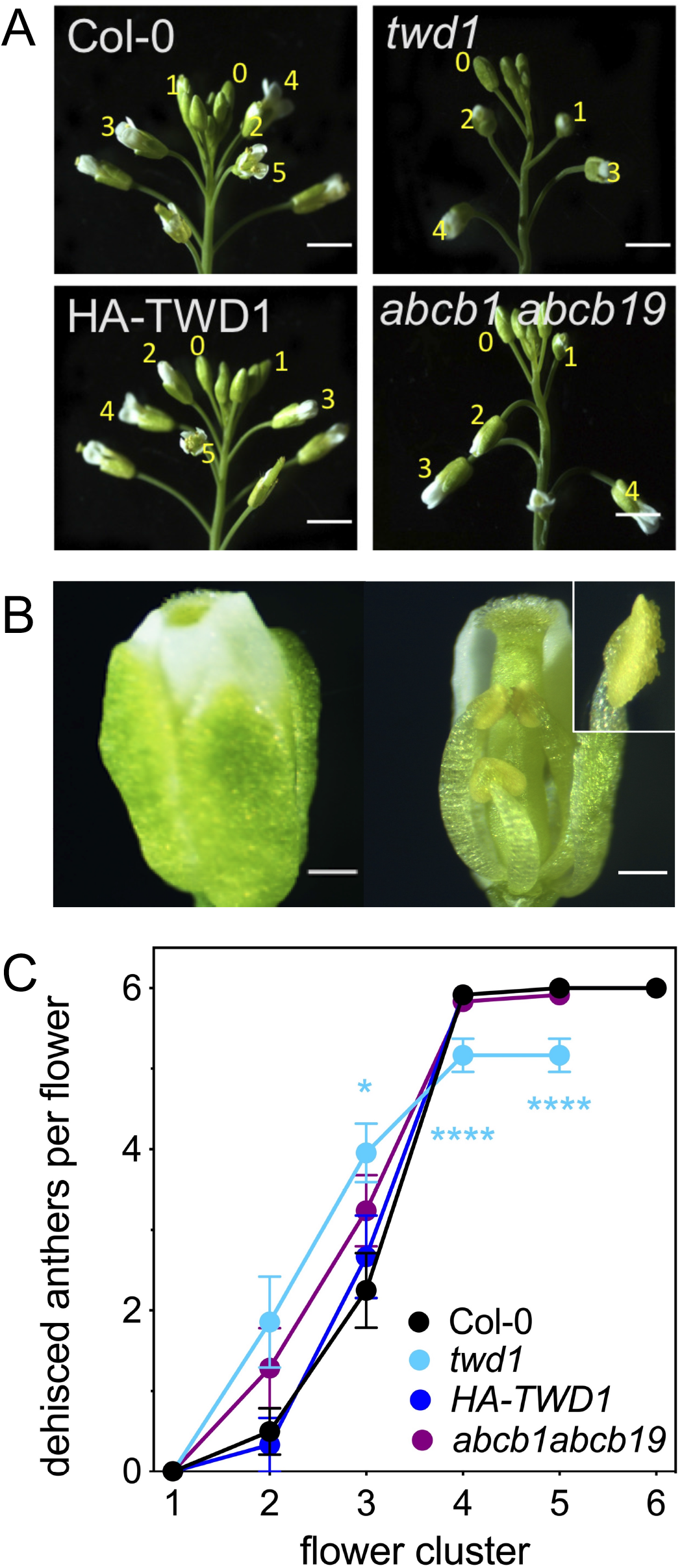
*twd1-3* anthers are early dehiscent. **A:** Flower clusters of wild-type (Col-0), *twd1 (twd1-3)*, HA-TWD1 (*35S:HA-TWD1* in *twd1-3)* and *abcb1 abcb19 (abcb1-100 abcb19-101)* used to quantitatively describe anther dehiscence. The number 5 indicates the end of floral stage 12. Bars, 250 μm. **B:** Partially dehiscent anthers (inset) are visible in *twd1-3* flowers at stage 12. Bars, 500 μm. **C:** The number of dehiscent anthers in wild-type (Col-0), *twd1 (twd1-3)*, HA-TWD1 (*35S:HA-TWD1* in *twd1-3)* and *abcb1 abcb19 (abcb1-100 abcb19-101)* flowers was determined at floral stage 13. Significant differences of mean values ± SEM (ordinary one-way ANOVA, Kruskal-Wallis test) to wild-type are indicated by asterisks (*, p<0.05; **, p<0.01; *** p<0.001).

In addition, *twd1* flowers show a reduced number of short stamens, which is surprisingly not found for the *abcb1abcb19* double mutant (Figure 1C). Auxin controls the early phase of floral organ primordia formation and morphogenesis by a combined action of local auxin biosynthesis, transport, and signaling that are all critical for stamen initiation (Cardarelli and Cecchetti, 2014; Cucinotta et al., 2020). However, ABCB transporters have no effect on the formation of stamen primordia, as both the single (*abcb1, abcb4, abcb19*) and double *abcb1 abcb19* mutant are not altered in the number of stamens, suggesting that the effect of *TWD1* on stamen primordia formation was not achieved through the so far best-characterized ABCB transporters, ABCB1, ABCB4 or ABCB19.

To further investigate flower development, we quantified the process of anther dehiscence within a flower cluster. The youngest flower with visible petals was labeled flower 1, the next elder flower was labeled flower 2, and so on (Figure 2A) (Jewell and Browse, 2016; Wei et al., 2018). We found that only a small number (8%) of *twd1* anthers at stage 12 showed significant early dehiscence (Figures 2B-C), which is exactly in line with reported phenotypes of *abcb1* but not of *abcb19* (Cecchetti *et al*., 2015). Interestingly, only 5 anthers dehisce in flower cluster 4 and 5 (Figures 2C), which might be connected to having less short-filament stamens in the mutant (see below).

Taken together, all the above results indicate that ABCB1 and ABCB19 provide coordinated auxin transport mechanisms during stamen development, and that TWD1 functions as a coordinator of these processes. The absence of significant differences in the single *abcb* mutants is probably due to partial redundancy in the function of ABCBs, which is well described for ABCB1 and ABCB19 (Geisler and Murphy, 2006), indicating that multiple ABCB-type auxin transport function interactively to regulate auxin transport during the late stages of stamen development. Despite the fact that PIN2 has also been reported to be localized in stamen filaments, and that loss-of-function mutation in *PIN2* result shorter filament in 30% of flowers (Kim et al., 2013), we did not find significant alterations in stamen development for *pin2* and *pin8* mutants (Supporting Information Fig. S1), suggesting that the described roles during PAT-mediated stamen development are eventually ABCB-specific.

### *TWD1* is expressed at both early and late stages of stamen development

To assess whether *TWD1* expression is consistent with the above-described phenotypes, longitudinal and transverse cross-sections of stamens at different developmental stages were analyzed by *in situ* hybridization. A highly specific anti-sense probe of 150 bp was designed that covers the TPR domain and a part of the calmodulin binding domains (aa 264-314) that are both thought to provide specificity to their binding partners by structural and not by sequence information (Geisler and Bailly, 2007). As a negative control, a sense probe revealed no signal at all at stages 8-10 (Supporting Information Fig. S2G). As shown in Figure 3A-C, the *TWD1* mRNA signal was detectable with stages 8 and 9 (premeiotic and meiotic stages (Bowman, 2012)) in the anther locules, mainly in the tapetum, middle layer, meiocytes (tetrads), and procambium. At early stage 10, the signal is weaker in the tapetum, middle layer and procambium but drastically enhanced in the tapetum and middle layer at late stage 10 (Figure 3D-E). At stage 11 (Figure 3F), *TWD1* mRNA is detectable in the procambium, in tapetum remnants (T*) and pollen grains. At stage 13 (Figure 3G), the TWD1 signal is no longer detectable, which is the same as previously reported for *ABCB1* (Cecchetti *et al*., 2015). The tissue-specific expression of *TWD1* was also consistent with the RNA-seq analysis results that were retrieved from a public-available database (http://travadb.org; Supporting Information Fig. S2A). Interestingly, also in these datasets the overall expression profile of *TWD1* resembles roughly *ABCB1* (but not of other *ABCB* genes) or *PIN2* and *PIN8* (Supplemental Figure 2A). These results verify the expression in the relevant stamen tissues and suggest that at least some of the functions of TWD1 during the regulation of anther development may correlate with ABCB1 actions.

**Fig. 3:**
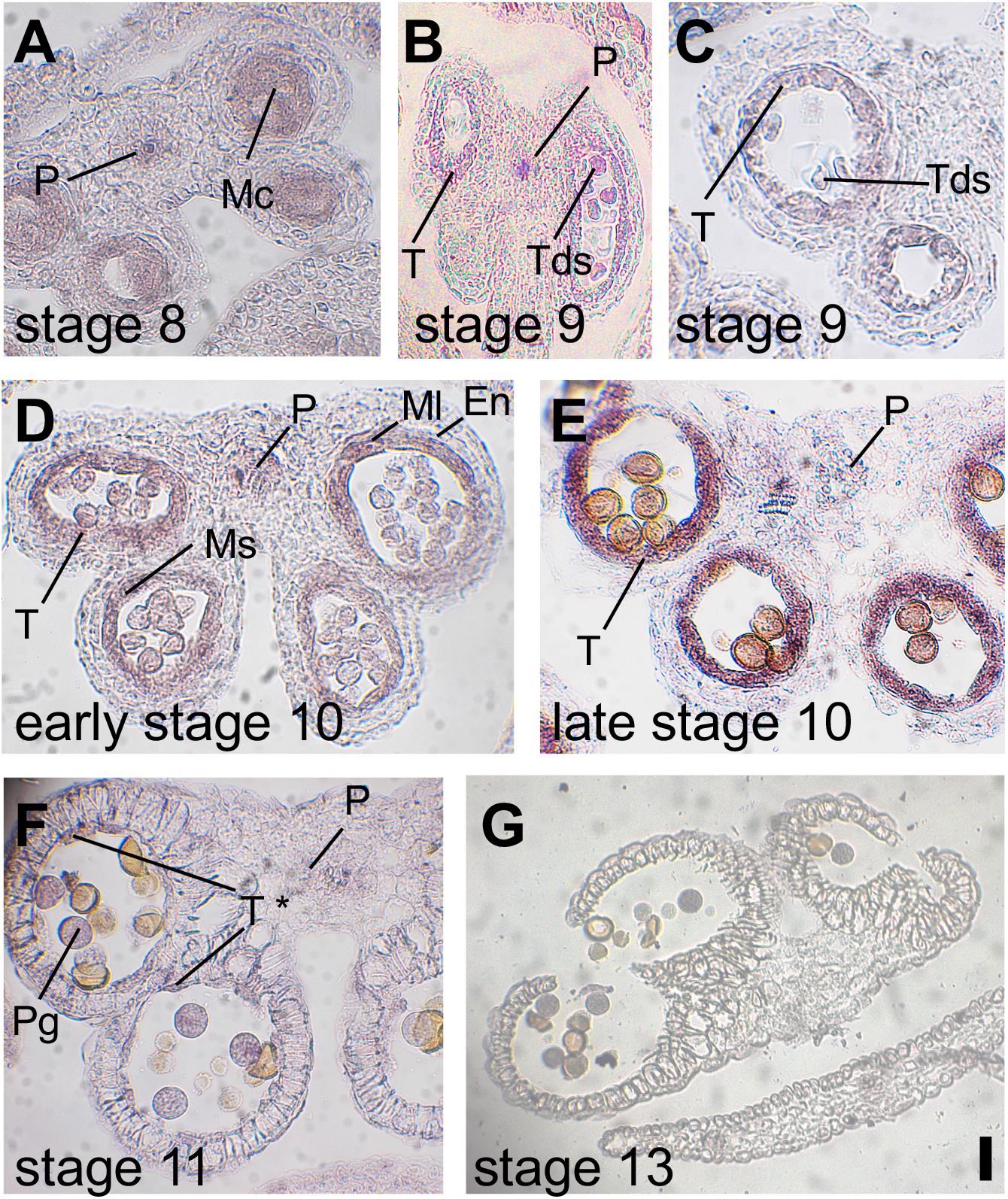
Expression of *TWD1* at early and late stages of stamen development. RNA *in situ* hybridization of *TWD1* using transverse sections of anthers at indicated stages (**A–G).** *TWD1* mRNA signal is localized in the anther locules, mainly in the tapetum (T), meiocytes (Mc), and the procambium (P) at meiotic and premeiotic stages (stages 8-9). The *TWD1* hybridization signal is weak in the tapetum (T), middle layer (ML), endothecium (En) and procambium at early stage 10 and is very strong in the tapetum at late stage 10. At stage 11, *TWD1* mRNA is detectable in the procambium, in tapetum remnants (T*) and in pollen grains (Pg). No signal is detectable at stage 13. Control hybridizations with sense probes for *TWD1* are shown in Figure S2. Mc, meiocytes; Ms, microspores; P, procambium; PG, pollen grains; SP, stamen primordium; T, tapetum; Tds, tetrads. Bars, 20 μm.

### Mislocalization of ABCB transporters in *twd1* impairs auxin transport in stamens

To gain further evidence for a correlation between ABCB-mediated auxin transport and TWD1 expression during stamen development, we imaged the expression of the auxin-responsive element DR5-GFP (Ottenschlager et al., 2003) in *twd1* (*twd1-3*; (Wang *et al*., 2013)) anthers and compared the GFP signals with those in the corresponding wild type (Col-0). DR5 expression had been previously detected in anthers from flower stages 9 to 12 (Cecchetti *et al*., 2008; Cecchetti *et al*., 2015; Cecchetti *et al*., 2017; Feng et al., 2006). The overall DR5-GFP expression pattern in *twd1* stamen anthers from flower stages 9 to12 was almost identical to that in the wild type, however, the expression level of DR5-GFP was significantly reduced (Figure 4A and Figure 4D) but fully restored in *twd1* complemented with *HA-TWD1* (not shown). In addition, the auxin maxima in the procambium were significantly reduced in the *twd1* mutant (Figure 4A), indicating that the irregular auxin accumulation caused by lack of auxin transport from the tapetum to the procambium is possibly associated with TWD1. To test this hypothesis, we carefully observed the details of the transit of DR5-GFP signals from the place of local auxin synthesis (tapetum) to the middle layer and the endothecium. The DR5-GFP signals in the endothecium of *twd1* was significantly decreased (Figure 4A and Figure 4D).

**Fig. 4:**
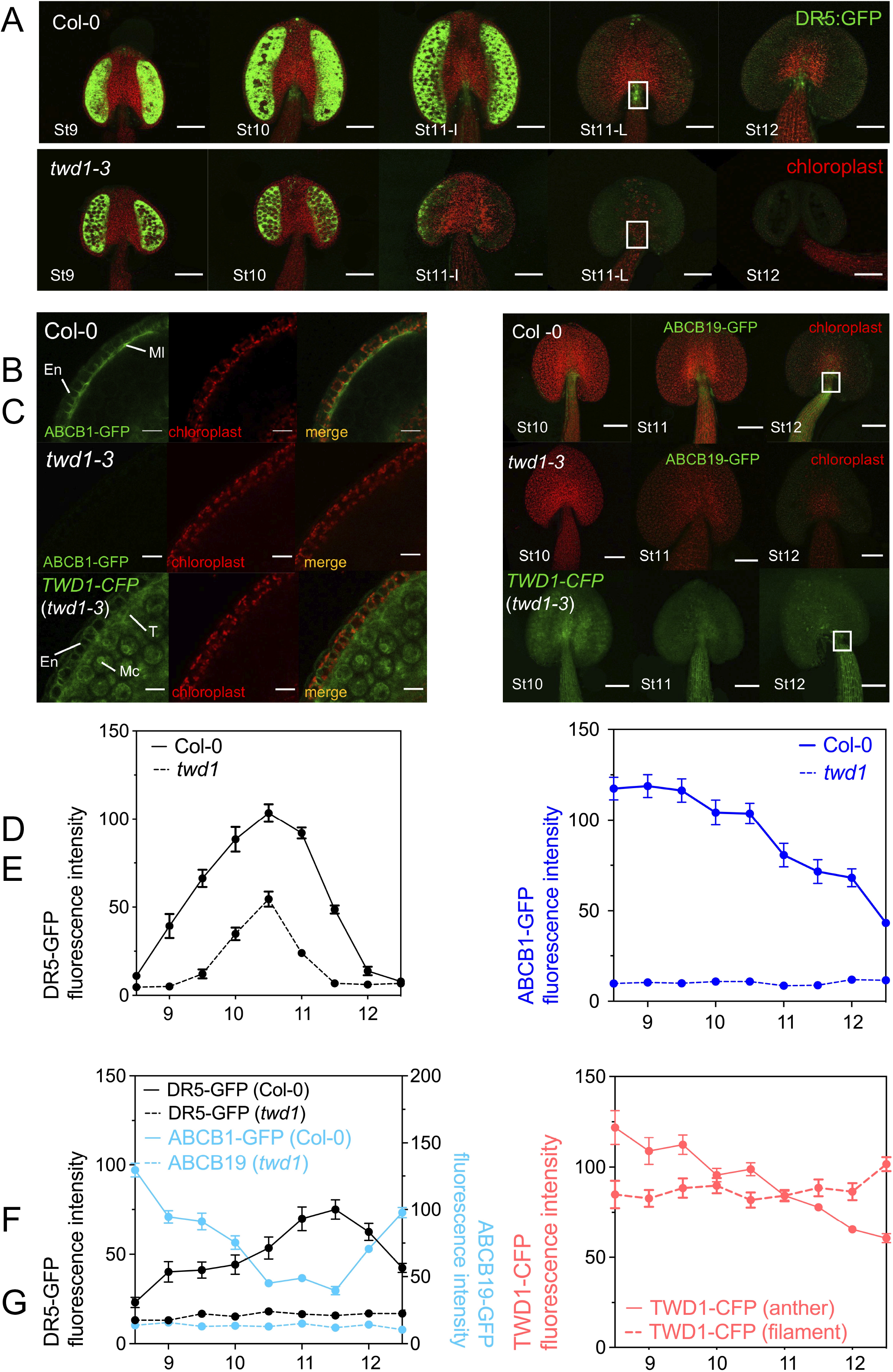
Effects of TWD1 on *Arabidopsis* long stamen development. **A:** Overall expression of the auxin responsive element, DR5:GFP, is significantly reduced in *twd1 (twd1-3)* during stamen development (floral stage 9-12). Note that the shift of maximum *DR5* expression in anthers to the procambium from early (St11I) to late initiation phase of stage 11 (St11L) is present in in wild-type (Col-0) but not in *twd1-3*. Green color indicates DR5-GFP signals, red color labels chloroplast autofluorescence; white boxes indicate the location of the procambium. Bars, 20 μm. **B:** Co-expression of ABCB1-GFP (imaged using *ABCB1:ABCB1-GFP)* and TWD1-CFP (images *usingTWD1:TWD1-CFP)* in the anther endothecium at early floral stage 10. Bars, 10 μm. **C:** Co-expression of ABCB19-GFP (imaged using *ABCB19:ABCB19-GFP)* and TWD1-CFP (images using *TWD1:TWD1-CFP* in *twd1-3)* in the anther endothecium at floral stages 10 to 12. White frames indicate the location of procambium in the same place as the white frames in A. Bars, 20 μm. **D**: Developmental time course of DR5-GFP expression in the anther of wild type (solid line) and *twd1 (twd1-3;* dotted line). **E:** Developmental time course of ABCB1-GFP expression (imaged using *ABCB1:ABCB1-GFP)* in the base of stamen filament of wild type (solid line) and *twd1 (twd1-3;* dotted line). **F:** Developmental time course of DR5-GFP (black lines) and ABCB19-GFP (light blue lines; imaged using *ABCB19:ABCB19-GFP)* expression in the filament tip region of wild type (solid line) and *twd1 (twd1-3;* dotted line). **G:** Developmental time course of TWD1-CFP expression (imaged using *TWD1:TWD1-CFP* in *twd1-3)* in the anther (red solid line) and filament (red dotted line). The expression profiles were collected from measuring 12-24 stamen from individual plants of each genotype.

In order to further support a putative regulatory role of TWD1 on ABCB1 and ABCB19 during this auxin transport process, the expression of TWD1, ABCB1 and ABCB19 was quantified on a protein level in stamen anthers and filaments. For that we used plants that expressed *TWD1::TWD1-CFP, ABCB1::ABCB1-GFP* and *ABCB1::ABCB1-GFP* in their mutant backgrounds (Mravec et al., 2008; Wu *et al*., 2010). Interestingly, ABCB1-GFP is not expressed in meiocytes and in the tapetum but can be detected in the middle layer of the anther at stage 10 and in the basal region of the filament at later stages (stages 10-11) during stamen development. ABCB1-GFP is then no longer concentrated at the base of the filament, but spreads throughout the stamen filament at stage 12 (Figure 4B and 4E; Supporting Information Movie S1). In addition, ABCB1-GFP was also highly expressed at the base of petals and sepals and at their receptacle junctions (Supporting Information Movie S1). In agreement with *in situ* data (Figure 3), TWD1 was predominantly expressed in the endothecium, the tapetum and the meiocytes at stage 10 (Figure 4B; Supporting Information Movie S3). Interestingly, both TWD1-CFP and ABCB1-GFP localized on the endothecium and middle layer plasma membrane (Figure 4B). Importantly, ABCB1-GFP was barely detectable in the *twd1* endothecium and middle layer, consistent with a significant decrease in DR5-GFP expression, suggesting that TWD1-mediated mislocalization of ABCB1 alters the auxin accumulation in the procambium at the inception of late stamen development.

Previous work had uncovered that *ABCB19* was predominantly expressed in the stamen filament (Noh *et al*., 2001) where it was found in both the epidermis and the pro-vasculature (Cecchetti *et al*., 2015). An investigation of ABCB19 on the protein level revealed that ABCB19-GFP expression was very strong in stamen filaments at flower stage 12 and was localized in both epidermal and provascular cells, while like for ABCB1 the ABCB19-GFP signal was lost in *twd1* stamen (Figure 4C and 4F; Supporting Information Movie S2). In addition, ABCB19-GFP was more concentrated below the DR5-GFP signal at the tip of the stamen filaments. Most interestingly, DR5-GFP expression increased with decreasing ABCB19-GFP expression during stamen development after floral stage 10 (Figure 4F). However, with the begin of the rapid filament elongation at stage 12, ABCB19-GFP increases again in the filaments, which is accompanied with a decrease in DR5-GFP fluorescence (Figure 4F). Excitingly, these events perfectly correlate with an increasing TWD1-CFP expression in filaments (but not in anthers) after floral stage 11 (Figure 4C and G).

Overall, these results support the idea that ABCB1 has a more stable, housekeeping-function, while ABCB19 expression fluctuates during stamen development. ABCB19 seems to function primarily in basipetal auxin transport from the stamen filament apical region to the basal side (Titapiwatanakun et al., 2009) as found for roots before (Blakeslee *et al*., 2007; Titapiwatanakun and Murphy, 2009). Thus, ABCB19 seems to affect stamen filament elongation by participating in auxin-mediated cell elongation and the specific high expression of ABCB19 at stage 12 might prepare for the rapid elongation of stamen filaments just before anthesis. Increasing ABCB19 and TWD1 expression correlate with the onset of rapid filament elongation, however, both ABCB1- and ABCB19-mediated auxin transport in anthers and filaments requires TWD1 action.

Next, we investigated whether significantly reduced auxin responses in the *twd1* stamen anthers this would have an effect on neighboring tissues, such as nectaries. In contrast to Wt (Aloni et al., 2006), the DR5-GFP signal in *twd1* nectaries completely disappeared in the lateral nectaries (Supporting Information Fig. S3B). This loss of auxin signaling in nectaries correlated with reduced starch accumulation in *twd1*, indicating reduced amounts of carbohydrate-rich nectar that is produced to attract visiting pollinators (Supporting Information Fig. S3C). Interestingly, this phenotype was not observed in the *abcb1 abcb19* double mutant. This dataset supports the concept that TWD1 affects nectary auxin homeostasis, however, in an action that is independent of ABCBs.

### TWD1 is important for male gametophyte development

It was previously assumed that more than 90% of the auxin synthesized by the tapetum would be stored in the pollen grains (Ljun et al., 2002). However, recent studies have shown that the early stages of pollen development require auxin produced in sporophytic microsporocytes rather than the tapetum (Yao et al., 2018). On the other side, it is assumed that the tapetum plays an active role in response to the secretion, breakdown, and synthesis of the essential pollen wall material, like callose (Xu et al., 2014), which contributes directly to microspore and pollen grain formation. TWD1 is localized in the tapetum and meiocytes (Figure 3A and Figure 4B) and TWD1-CFP was detectable since the tetrade stage (Supporting Information Fig. S4B). Asynchronous development in the locules of the same anther was frequently observed in *abcb1 abcb19* around the end of meiosis (Supporting Information Fig. S4C; (Cecchetti *et al*., 2015)).

Here we report that both pollen viability and pollen number was significantly reduced in the *twd1* and the *abcb1abcb19* double mutant but not in investigated *pin* mutants (Gao et al., 2021); Figure 5A-B). DAPI staining of pollen revealed aborted, misshaped and multi-nucleus pollen in *twd1* and *abcb1 abcb19* (Figure 5C; Supporting Information Fig. S4C). Furthermore, the *twd1* microspores failed to form the cell plate (Figure 5D), indicating defects in pollen mitosis. The TWD1 expression in the tapetum is consistent with the reduced pollen viability phenotype of *twd1*. Based on the tapetum localization of TWD1, it might be speculated that TWD1 may also be involved in cell wall establishment during pollen maturation.

**Fig. 5:**
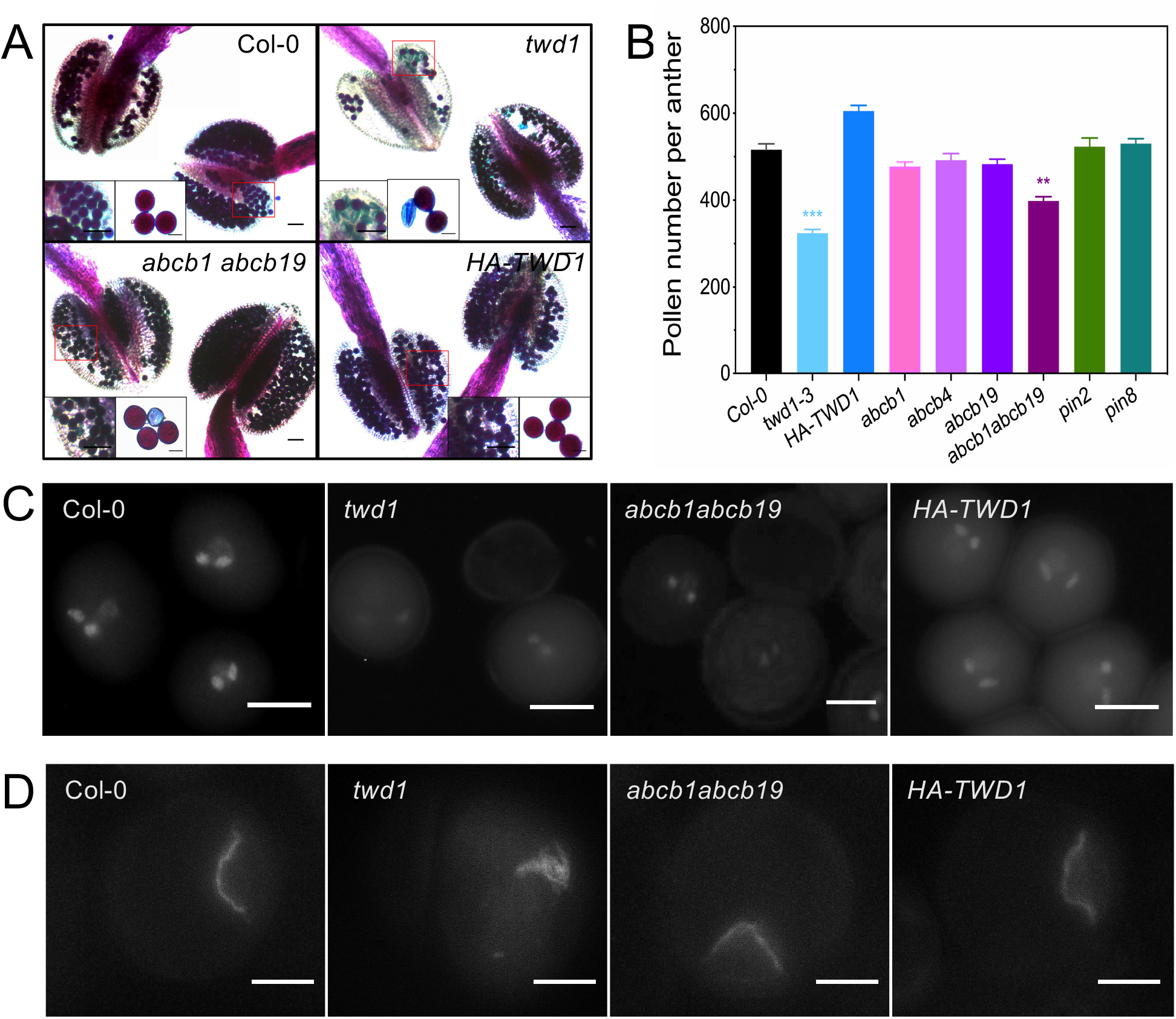
*twd1* pollen grains are shriveled and inviable. **A:** *twd1 (twd1-3)* and *abcb1 abcb19 (abcb1-100 abcb19-101)* anthers contain viable and dead pollen grains as indicated by their abnormal shapes and green coloration in Alexander stains. Bars, 50 μm. **B:** *twd1 (twd1-3)* and *abcb1 abcb19 (abcb1-100 abcb19-101)* anthers contain less pollen grains. Significant differences of mean values ± SEM (ordinary one-way ANOVA, Kruskal-Wallis test) to wild-type are indicated by asterisks (*, p<0,05; **, p<0,01, *** p < 0,001). One hundred anthers from four individual plants of each genotype were analyzed before anther dehiscence shown in A. **C:** *twd1 (twd1-3)* and *abcb1 abcb19 (abcb1-100 abcb19-101)* pollen reveal aberrant or absent nuclei revealed by DAPI stains. Bars, 10 μm. **D:** Callose accumulation labeled by aniline blue in wild type (Col-0), *twd1 (twd1-3), abcb1 abcb19 (abcb1-100 abcb19-101)* and HA-TWD1 (*35S:HA-TWD1* in *twd1-3)*. Note that in *twd1* but not *abcb1 abcb19*, callose accumulated as a large aggregate at the cell cortex indicating that the cell plate failed to be formed in defective microspores. Bars, 10 μm.

As a consequence, we observed a high frequency of sterility in *twd1* and *abcb1 abcb19* flowers that produced siliques with reduced lengths and numbers of seeds per silique (Figure 6A-C). Mature and dry siliques from *twd1* and *abcb1 abcb19* were found to have ~20 % empty spaces compared with wild type (~3% empty spaces; Figure 6D-E). While silique length and seed number were also slightly but significantly affected in *abcb1* and *abcb19* single (but not in investigated *pin*) mutants, this was not the case for seed abortion indicating a different degree of functional redundancy between these isoforms during these developmental processes. As previously described, the mature *abcb1 abcb19* gynoecium appeared morphologically similar to that of the wild type (Larsson et al., 2014), suggesting that the sterility of mutants was due to the failure of the stamen filament elongation to position the anther above the stigma at anthesis, presumably in *twd1* as well.

**Fig. 6:**
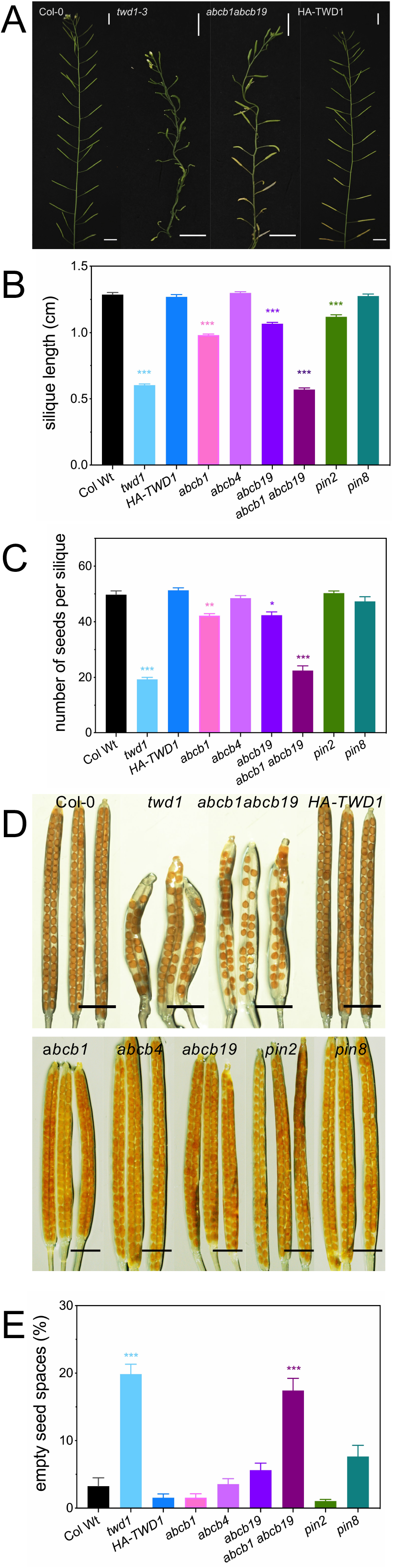
*twd1-3* siliques have aborted seeds. **A:** Flowers of wild-type (Col-0), *twd1 (twd1-3)*, HA-TWD1 (*35S:HA-TWD1* in *twd1-3)* and *abcb1 abcb19 (abcb1-100 and abcb19-101)*. Bars, 1 cm **B-C:** *twd1 (twd1-3)* and *abcb1 abcb19 (abcb1-100 abcb19-101)* siliques are shorter (**B**) and contain a reduced number of seeds per silique (**C**). Significant differences of mean values ± SE (ordinary one-way ANOVA, Kruskal-Wallis test) to wild-type are indicated by asterisks (*, p<0,05; **, p<0,01, *** p < 0,001). One hundred flowers from four individual plants of each genotype were analyzed before anther dehiscence shown in A. **D-E:** Immature siliques from indicated lines (**D**). Bars, 200 μm. Quantification of empty spaces in mature siliques from indicated lines indicating seed abortion. Significant differences of mean values ± SE (ordinary one-way ANOVA, Kruskal-Wallis test) to wild-type are indicated by asterisks (*, p<0,05; **, p<0,01, *** p < 0,001). One hundred flowers from four individual plants of each genotype were analyzed.

Overall, our data indicate that *TWD1* mutation results in a reduced auxin flow in stamen filaments leading to defects in three processes of late stamen development: stamen elongation, anther dehiscence and pollen maturation. Moreover, altered PAT in filaments is responsible for altered auxin responses in neighbor tissues (nectaries) and results in defective pollen development and seed sterility, which qualifies TWD1 as key regulator of male gametophyte development. In most of these processes, TWD1 seems to perceive its action by a differential activation of ABCB-mediated auxin transport (mainly: ABCB1). In support of this, TWD1 and ABCB1 share widely overlapping *in situ* pattern (Figure 3) and developmental phenotypes (Cecchetti *et al*., 2015).

## Discussion

### TWD1 functions as a key player in Arabidopsis stamen development

Auxin biology has focused over the last decades mainly on root development, which is mainly based on the fact that imaging of auxin transporters and auxin reporters is facilitated in the transparent root tissues. Despite clear evidence that local auxin accumulation, polar transport and auxin signaling are all likewise critical for floral organ initiation (Cardarelli and Cecchetti, 2014; Cucinotta *et al*., 2020), much less is known about the molecular key players that regulate auxin-controlled stamen development.

In this study, we have analyzed stamen development of the *twd1* mutant that on a first view in comparison to its vegetative tissues looks less effected. This is also reflected by reduced but reasonable transformation rates during floral dipping of *twd1* alleles that allowed for genetic access. However, a careful analysis revealed strong defects in flower size and shape, in the physical position of the anthers and in long stamen elongation for the *twd1* mutant (Figures 1-2). With no surprise, like previously described for vegetative organs, most of the described defects in *twd1* are shared with the *abcb1 abcb19* mutant (Wang *et al*., 2013), which can be attributed to the fact that TWD1 as a described ABCB chaperon is a positive regulator of ABCB1 and ABCB19 action.

Most likely as a consequence of altered auxin delivery (Figure 4; for details see below), both viability and number of pollens was significantly reduced in *twd1* (and *abcb1abcb19*) that were often aborted, misshaped and contained multiple nuclei (Figure 5), indicating defects in pollen mitosis. This resulted in a high frequency of sterility in *twd1* (and *abcb1 abcb19*) flowers producing less seeds per silique (Figure 7) reducing overall plant fitness.

A surprising finding was that short stamen development was also under control of TWD1, however, apparently independent of ABCB1,4,19 (and PIN2,8), which is indicated by the finding that short stamen number was not significantly different from wild-type in single or double *ABCB* mutants (Figure 1C). In agreement, also early anther dehiscence (Figure 2; (Cecchetti *et al*., 2015)) and starch accumulation in lateral nectaries (Supporting Information Fig. S3) was dependent on TWD1 but not on investigated ABCBs. Interestingly, PIN6, a member of the ER-localized, so-called short-loop PINs, specifically localized in the nectaries and young stamens, is positively correlated with short stamen formation and total nectar production (Bender et al., 2013). Currently, it is unclear whether PIN6 and TWD1 are directly related or similarly regulated during stamen formation and nectary maturation. A plausible explanation might be that TWD1 either also regulates PIN6 auxin transport activity by an action that is dependent on its chaperone activity and thus requires direct protein-protein interaction or it involves the auxin-actin circuit (Tan et al., 2020; Zhu *et al*., 2016).

### TWD1 functionally interacts with ABCBs in regulating auxin accumulation and transport during late stamen development

Previously, the role of ABCB1 and 19 in stamen development was mainly deduced from gene expression profiling, which can have many pitfalls because mRNA levels do not always translate directly into protein expression. Here, we have quantified the expression profiles of ABCB1 and 19 and its regulatory chaperon, TWD1, on the protein level during flower development by using established fluorescent protein reporter lines in their mutant backgrounds (Mravec *et al*., 2008; Wu *et al*., 2010). Both TWD1 and ABCB1 are localized on the middle layer, endothecium plasma membrane, and procambium, while ABCB1-GFP was barely detectable in *twd1* stamen (Figure 3 and Figure 4B). Auxin maxima in the procambium were altered in *twd1* mutant (Figure 4A), suggesting a polar auxin transport from the anther to the procambium. Our results suggest that both TWD1 and ABCB1 localize to at the plasma membrane of the middle layer and the endothecium where they co-function in auxin transport from the middle-layer to the procambium. Their absence results in an inhibition of auxin transport from the middle layer to the outer layer and the reduction of DR5-GFP signal in the *twd1* procambium and endothecium (Figure 4A) is a consequence of this inhibition. Combining our results with previous findings (Cecchetti *et al*., 2015) suggests that the middle layer acts as a “dam of the auxin river”, blocking upstream auxin flow from the tapetum to the endothecium and procambium. In this comparison, ABCB1 functions as the gate of the dam and regulates the auxin level of the middle layer dam. In contrast, the endothecium and procambium would be the downstream “hydrological station”. This is supported by reduced auxin levels downstream in the absence of TWD1, which are most likely caused by immature ABCB1 at the plasma membrane that is unable to fulfil its function as a gate-opening switch in the middle layer dam the absence of the TWD1 chaperone.

Auxin accumulation in the procambium peaks at the end of stage S11 in the junction region between filament and anther (Cecchetti *et al*., 2008), followed by a rapid increase in the expression of ABCB19-GFP, that concentrated just below the maximum auxin position at the tip of the stamen filaments (Figure 4C and Figure 4F). Subsequently, DR5-GFP and ABCB19-GFP expression correlated negatively late stamen development (stage 12; Figure 4F), obviously preparing for the rapid elongation of stamen filaments just before anthesis. Previously, ABCB19 was considered primarily to function in basipetal auxin transport from the apical region of the stamen filament to the basal side (Blakeslee *et al*., 2007; Titapiwatanakun and Murphy, 2009). Our observations suggest that ABCB19 localized in the provasculature cells is responsible for the basipetal auxin transport from apical to basal sides of the filament, in synergy with ABCB1 localized in the basal region of stamen filaments (Supporting Information Movie S1).

Based on the overall stable and fluctuating expression profiles for ABCB1 and ABCB19, respectively, and correlating DR5-GFP levels, we predict an overall housekeeping function for ABCB1 during earlier stages, while ABCB19 seems to be responsible for the key event of rapid elongation at later stages of stamen development together with ABCB1 (Figure 4). Despite the fact that TWD1 is also a functional ABCB4 chaperon (Wang *et al*., 2013; Wu *et al*., 2010), we could not find any prominent role for ABCB4 in stamen development, which is obviously also in line with its absence in flowers and its prominent root expression profile (Supporting Information Fig. S2C).

Altogether, our results allow us to propose a model summarized in Supporting Information Fig. S5, in that polar auxin transport from the middle layer to the outer procambium by ABCB1 is mainly responsible for the formation of an auxin maximum in the latter at the inception of late development. From the procambium, auxin is then transported basipetally towards the basal side of the filament as previously proposed (Cecchetti *et al*., 2017) mainly through ABCB19 causing rapid stamen filament elongation just before anthesis. A possible additional contribution of ABCB1 to filament elongation is also suggested by the expression of ABCB1 at stage 12 in the basal part of the anther. TWD1 differentially activates ABCB1 and 19 during auxin transport providing thus local auxin accumulation during stamen development. This dynamics auxin distribution from the inner tissue to the outer tissues would guide filament elongation, thus ultimately supporting pollen development and finally fertilization of the ovules by the pollen.

## Supplementary data are available at *JXB* online

**Fig. S1: TWD1 loss-of-function flowers have a defect in stamen development**

**Fig. S2: Gene expression profiles of TWD1 and selected auxin transporters and in situ hybridization controls**

**Fig. S3: TWD1 is essential for male gametophyte development**

**Fig. S4: TWD1 is required for proper auxin responses in the flower nectaries.**

**Fig. S5: Hypothetical model showing auxin accumulation and ABCB-mediated transport during stamen development**

**Movie S1: Expression of ABCB1-GFP (ABCB1:ABCB1-GFP) in stamen at floral stage 10 and 12.**

**Movie S2: Expression of ABCB19-GFP (ABCB19:ABCB19-GFP) in stamen at floral stage 10 and 12.**

**Movie S3: Expression of TWD1-GFP (TWD1: TWD1-GFP) in stamen at floral stage 10 and 12.**

## Acknowledgements

This work was supported by Swiss National Funds (project 31003A_165877 and 310030_197563) to M.G.

## Author contribution

M.G and J.L. designed the project and J.L. performed most experiments except the *TWD1 in situ* hybridization, which was performed by R.G. R.G. also assisted in the flower analyses. J.L. and R.G were supervised by M.G. and M.C., respectively. Data were analyzed by J.L., R.G., M.C. and M.G. and J.L. and M.G. wrote the manuscript; all authors commented on the manuscript.

